# Biological motion perception in the theoretical framework of perceptual decision-making: An event-related potential study

**DOI:** 10.1101/2023.09.16.558064

**Authors:** Osman Çağrı Oğuz, Berfin Aydın, Burcu A. Urgen

## Abstract

Biological motion perception plays a critical role in various decisions in daily life for humans. Failure to decide accordingly in such a perceptual task could have life-threatening consequences. Neurophysiological studies on non-human primates and computational modeling studies suggest two processes that mediate perceptual decision-making. One of these signals is associated with the accumulation of sensory evidence and the other with response selection. Recent EEG studies with human participants have introduced an event-related potential called Centroparietal Positive Potential (CPP) as a neural marker aligned with the sensory evidence accumulation while effectively distinguishing it from motor-related event-related potential, namely lateralized readiness potential (LRP). The present study aims to investigate the neural mechanisms of biological motion perception in the theoretical framework of perceptual decision-making, which has been overlooked before. More specifically, we examine whether CPP would track the coherence of the biological motion stimuli and could be distinguished from the LRP signal. We recorded EEG from human participants while they were engaged in a direction discrimination task of a point-light walker whose coherence was manipulated by embedding it in various levels of noise. Our behavioral findings revealed shorter reaction times and reduced miss rates as the coherence of the target stimuli increased. In addition, CPP tracked the coherence of the biological motion stimuli with a tendency to reach a common level during the response, albeit with a later onset than the previously reported results in random-dot motion paradigms. Furthermore, CPP was distinguished from the LRP signal based on its temporal profile. Overall, our results suggest that the mechanisms underlying perceptual decision-making generalize to more complex and socially significant stimuli like biological motion. We discuss the implications of these findings for advancing computational models of biological motion perception and perceptual decision-making.

## INTRODUCTION

One of the fundamental roles of the human visual system lies in its capacity to perceive and discern animate movements within its surroundings. Notably, perception of biological motion influences various judgments, including the recognition of living entities, agency attribution, responding to and interacting with those agents in the environment, and shaping broader social cognition. The human visual system remarkably and efficiently executes this function (Blake & Shiffrar, 2007; Pavlova, 2012; Rutherford & Kuhlmeier, 2013; Johnson & Shiffrar, 2013; Brooks et al., 2007). To illustrate, envision a scenario of driving a car when suddenly confronted with a pedestrian or a dog. Our visual system promptly detects the motion characteristic of a living being, which results in an immediate deceleration or halt. Without this essential ability, the consequences could become life-threatening. This highlights the significance of delving into biological motion perception within the realm of perceptual decision-making.

Perceptual decision-making and its neural mechanisms have been widely researched in systems and cognitive neuroscience by utilizing techniques ranging from invasive neurophysiology in non-human primates to non-invasive neuroimaging in humans and computational modeling. A frequently employed paradigm for investigating perceptual decision-making in empirical studies is the random-dot paradigm. This paradigm involves the display of randomly moving dots, some of which are moving in the same direction (Newsome et al., 1989). The dots moving in the same direction are called as *signal dots*, while the remaining dots moving in random directions are referred to as *noise dots*. By adjusting the proportion of *signal dots*, researchers can manipulate the coherence of the motion stimuli. This paradigm is thought to effectively simulate real-world scenarios where stimuli are often imprecise and contaminated by noise. Historically, it has been primarily used with non-human primates while recording the neural firing rate in different brain areas to reveal the underlying mechanisms of perceptual decision-making (Horwitz & Newsome, 1999; Kim & Shadlen,1999; Shadlen & Newsome,1996). An influential neurophysiology study conducted by Shadlen & Newsome (2001) examined neural firing rates time-locked to the stimulus or response onset while monkeys reported the direction of movement with eye movements (saccades). The behavioral results demonstrate that increased coherence of the motion stimuli is associated with shorter mean reaction times and higher accuracies. More importantly, for the neurophysiological findings, increased stimulus coherence correlated well with elevated neural firing rates in the lateral intraparietal area (LIP). This pattern stands in contrast to findings from the middle temporal area (MT), a brain region typically associated with motion perception. Furthermore, researchers observed that activity from LIP neurons reached a uniform activity level during the response, irrespective of stimulus coherence. These findings collectively suggest that LIP is not solely involved in motion perception but also plays a crucial role in perceptual decision-making (Shadlen & Newsome, 2001; Gold & Shadlen, 2007).

The insights gained from neurophysiological investigations in non-human primates have been integrated into various computational models, with one prominent example being the drift-diffusion model (Smith & Ratcliff, 2004; Ratcliff & McKoon, 2008). The model consists of two main components: the accumulation of sensory evidence and the decision boundary. Essentially, the drift-diffusion model proposes that during perceptual decision-making, our sensory system initially collects and accumulates evidence over time, ultimately surpassing a predetermined threshold for the decision to be reached, namely the decision boundary. To illustrate, in the context of random-dot motion experiments, the relationship between the time required for response and maximum neural firing rates aligns with the stimuli coherence (Bach et al., 2011), indicating the progressive accumulation of sensory evidence.

On the other hand, when investigating perceptual decision-making in humans, researchers predominantly utilized non-invasive techniques, one of which is high-temporal-resolution EEG (Kelly and O’Connell, 2013, 2015; Twomey et al., 2015). These studies have employed event-related brain potentials alongside behavioral measures such as reaction times and miss rates as the key dependent variables. In their seminal work, Kelly and O’Connell (2013) manipulated the coherence level of random dot stimuli, akin to previous non-human primate research, and instructed participants to report the direction of movement. However, unlike previous neurophysiology studies, the target motion stimuli were preceded by 0% coherent dots, namely *scrambled dots*. This continuous-stimulation design allowed for a seamless transition from the scrambled stimulus to the target stimulus, preventing any immediate visual evoked potentials upon motion stimuli presentation. The results revealed that stimuli coherence increase is associated with reduced miss rates and mean response times, consistent with prior findings. More importantly, the study revealed an event-related potential, Centro-Parietal Positivity (CPP), characterized by a positive distribution over centroparietal electrodes and emerging between 200-800 ms following the target presentation. CPP was observed to evolve over time in accordance with sensory evidence strength. More specifically, shorter peak latency and larger amplitude were observed in CPP, following the increase in target stimulus coherence. Furthermore, response-locked analyses indicated that CPP signals across different coherence levels converged at the response time. Therefore, CPP is thought to reflect both the sensory evidence accumulation process and the decision boundary postulated by the drift-diffusion model.

Numerous other studies have demonstrated complementary outcomes to CPP-related work on perceptual decision-making, encompassing various discrimination tasks and stimuli. In the visual domain, Philiastides et al. (2006) utilized a face-car categorization task and revealed an event-related component approximately 300 ms after stimulus onset that is thought to reflect the sensory evidence accumulation process. In the auditory domain, Kaiser et al. (2006) utilized an auditory experiment where the participants had to decide on the identity or location of syllables using MEG. The results demonstrated a similar ERP component, with more pronounced amplitude occurring at approximately 280–430 ms, for easier decisions than difficult ones, regardless of changes in identity or acoustics (Kaiser et al., 2006). Within the somatosensory domain, Herding et al. (2019) employed sequentially presented vibrotactile stimuli and asked participants to compare the frequencies. The findings demonstrated that the CPP signal aligns with subjectively perceived decision evidence (Herding et al., 2019). Collectively, these studies further suggest that CPP systematically builds up depending on the evidence strength, irrespective of sensory modalities (visual, auditory, or somatosensory), the manipulations targeting signal-to-noise ratio of the targets (e.g., intensity or volume changes), or the motor demands of the task including button presses (Kelly & O’Connell, 2015).

Despite the progress in the field, previous investigations into perceptual decision-making in the visual domain have predominantly employed relatively simple or static stimuli (e.g., random dot motion, Gabor patches, varying-sized circles, faces, and cars) (Kelly & O’Connell, 2013; Heekeren et al., 2008; de Lange et al., 2010; Cheadle et al., 2014; Gorea et al., 2014). However, whether the underlying processes for these more straightforward tasks generalize to more complex and socially significant situations involving motion, such as biological motion perception, remains unknown.

Point-light displays (PLDs) have emerged as fundamental stimuli for investigating biological motion perception. These displays consist of dots representing the joints of a moving actor (Johansson, 1973). Although they lack some fine details typically present in the natural motion of biological organisms, such as color, texture, and depth information, PLDs still offer a valuable tool for studying the visual system’s ability to detect and process animate movement (Blake & Shiffrar, 2007; Troje & Westhoff, 2006). One notable advantage of point-light displays is their flexibility for manipulation. Researchers frequently employ scrambled motion as a control stimulus by randomizing the initial positions of the dots in PLDs (Blake, 1993; Saygin et al., 2004; Lapenta et al., 2017). Moreover, methods have been developed to manipulate the coherence of PLDs by introducing a certain number of dots moving randomly, a task that would be challenging with natural movies, but which becomes feasible through PLDs (Decatoire et al., 2019). This approach has been widely adopted in behavioral, TMS, and fMRI studies, enabling researchers to easily manipulate the signal-to-noise ratio of biological motion stimuli (van Kemenade et al., 2012; Miller & Saygin, 2013; Gilaie-Dotan et al., 2013).

Given the evidence obtained from diverse stimuli and modalities as previously outlined, one could anticipate that mechanisms underlying perceptual decision-making extend to the biological motion processing. However, to the best of our knowledge, biological motion perception has yet to be explicitly studied within the theoretical framework of perceptual decision-making, representing a promising area of research. We believe such an investigation could broaden the scope of perceptual decision-making studies while also holding significant potential for enhancing our understanding of the neural mechanisms of biological motion perception, facilitating the development of more powerful computational models, and situating it within the broader context of visual perception.

This present study investigates the neural mechanisms of human biological motion perception in the theoretical framework of perceptual decision-making. To this end, we recorded behavior and EEG activity during a direction discrimination task of a point light walker stimulus embedded in various noise levels. We hypothesized that the mean reaction time and miss rate would change as a function of the coherence. More specifically, we predicted that increasing the noise level (i.e., decreasing the coherence) would increase the mean reaction time and the miss rate. Regarding ERPs, we expected that CPP would track the coherence level of the biological motion stimuli. However, we expected a potentially *later* onset for CPP than in previous work since the perception of biological motion may require more time to process than random dot motion stimuli due to its complexity. In addition, we also examined LRP as an index of motor preparation (Gratton et al.,1988) and aimed to distinguish it from CPP, following the previous work (Kelly & O’Connell, 2013).

## METHODS AND MATERIALS

### Participants

A total of 16 participants (8 female, mean age = 23.18, age range [18 40]) with normal or corrected-to-normal vision participated in the study. None of the participants had a neurological or psychiatric disorder and use any psychiatric medication. The Human Research Ethics Board of Bilkent University approved the study. Participants were informed about the study procedures and signed a consent form before participation. In the pre-screening form, all participants reported at least 6 hours of sleep.

### Stimuli, Experimental Design, and Procedure

The experiment utilized point-light displays for the biological motion stimuli to investigate human perceptual decision-making with EEG, renowned for their efficacy in capturing biological motion dynamics. The biological motion stimuli in the task portrayed a human walking action, featuring 13 dots representing the joints on both sides of the body and the head (with an overall size of 6.5 degrees). The walking direction could be either to the right or left while the stimuli were displayed from the side view of the corresponding direction.

For the scrambled motion stimuli, the motion vector of the dots mirrored that of the meaningful biological motion stimuli. However, instead of moving in a coherent direction to simulate human walking, the dots appeared to move randomly since their starting coordinates were randomized. Similar to the biological motion stimuli, varying degrees of noise dots were introduced into the scrambled motion displays. To maintain uniformity within each trial, the number of noise dots introduced for the scrambled motion stimuli matched those introduced for the subsequent biomotion stimuli.

For coherence manipulation, the biological motion stimuli were embedded in different levels of noise. Specifically, different quantities of dots were introduced into the display, effectively determining the trial’s noise level. These noise levels were determined through a pilot study, ensuring that manipulation of difficulty spanned four discernible levels. As a result, manipulation of difficulty was accomplished across four distinct noise levels: 10, 20, 35, 55.

Each of the four noise levels featured 120 trials, resulting in 480 trials in total for the whole experiment. The experiment was structured into four blocks, each consisting of 120 trials of varying levels of noisy stimuli. The counterbalancing mechanism ensured trial uniformity, encompassing both the direction of biological motion stimuli and the noise levels.

A preliminary practice task was administered for participants’ familiarization with the task. Participants were also instructed to fixate on the center - where the fixation point was displayed - while performing the task. Participants were viewing the screen with 57 cm distance. The participant’s task was to report the walking direction during the target interval using bimanual key presses. Each trial started with the presentation of a fixation point for 1 second at the trial’s onset, placed at the center of the screen to direct the participant’s attention to the designated area. Then, it was followed by the scrambled motion display. The scrambled motion stimuli were displayed at varying durations (3, 5, or 8 secs) to prevent the predictability of the target stimulus. After the scrambled motion, the target biological motion stimulus was displayed for a maximum of 2 seconds (See Figure 1). The transition from scrambled to biological motion stimuli was executed seamlessly to eliminate potential immediate visual evoked potentials in the EEG data.

**Figure 1.**
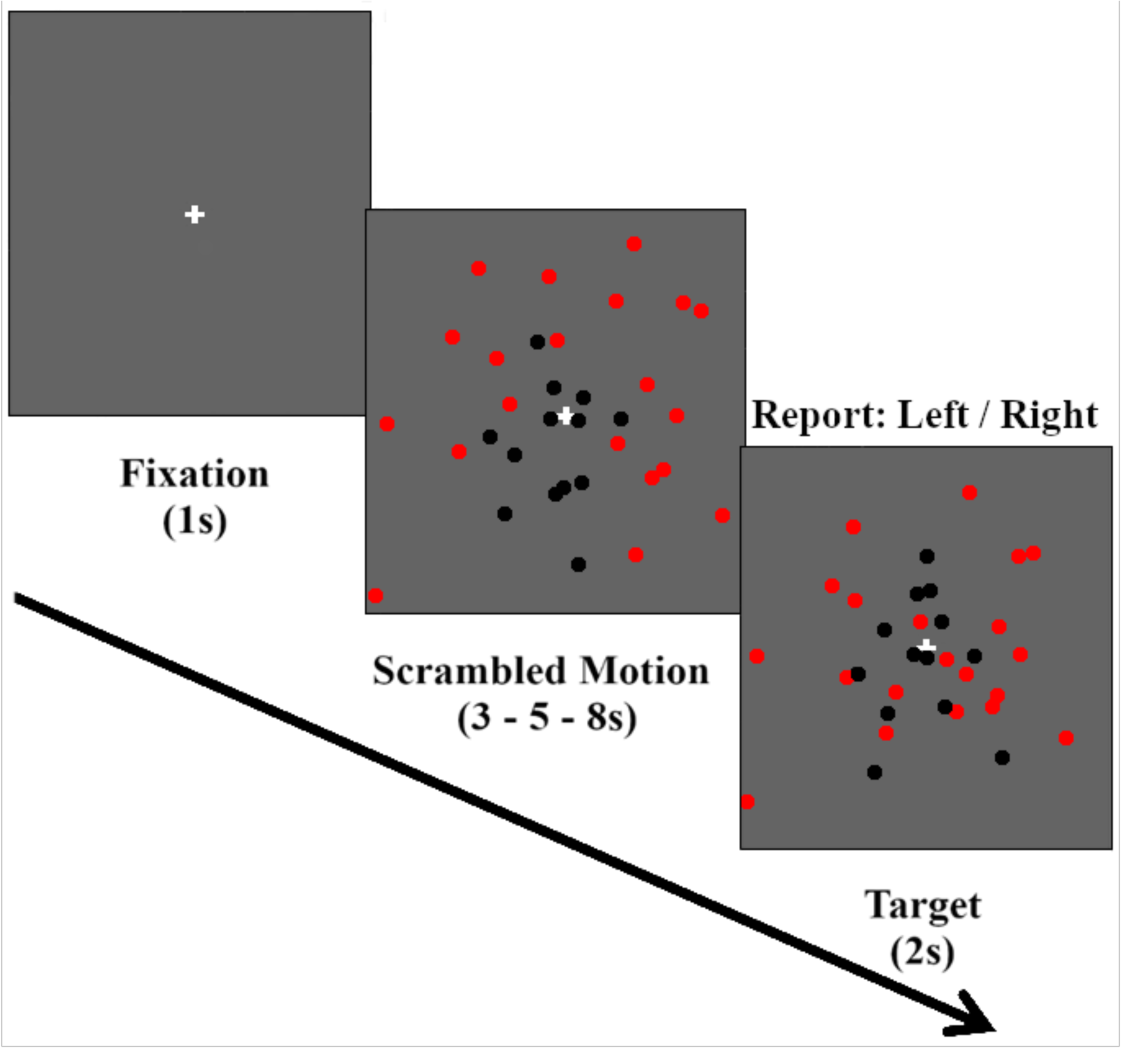
Experimental Setup. The example trial is illustrated with noise level 20 and the walking direction to the left. The red dots are depicted to highlight the noisy dots and all of the dots are in larger sizes than the actual stimuli for display purposes.

### Data Collection and Analysis

#### Behavioral

The mean response time and miss rate for the direction discrimination task were recorded and analyzed for each noise level. Response time, which is defined as the period that spans the onset of the biological motion stimulus until the participant gives a response, was taken for the correct trials. The miss rate indicates the proportion of trials where participants failed to respond during the target stimulus presentation. This measure provides information on the participant’s ability to detect the target and respond on time. One-way repeated measures analysis of variance (ANOVA) was applied to examine how different noise levels affected the mean response time and miss rate. Our hypothesis posited that as noise levels increased, response time and miss rates would also increase.

#### EEG Recording and Pre-processing

64-channel EEG (Brain Products, GmbH, Germany) was recorded at 512 Hz using sintered Ag/AgCl passive electrodes whose placements were according to the international 10/20 system. The electrode AFz was used as ground and FCz was used as reference. Electrode offset was kept below 25 k ohm. Pre-processing was done using Matlab and the EEGLAB toolbox (Delorme & Makeig, 2004). The analysis focused on 63 EEG channels after excluding the ECG channel. Data underwent a 1 Hz high-pass and 50 Hz low-pass filter, followed by re-referencing to the average of all electrodes. Eye-related components were removed using Independent Component Analysis (ICA), specifically by the binica algorithm. Data were then epoched for stimulus-locked and response-locked analysis. Epoch time intervals were determined based on the mean response times instead of minimum response times for accurate trials for each participant due to the insufficiency of the latter to capture the entire decision process. The mean response time was approximately 830 ms. Thus, our analyses incorporated 800 ms post-target and pre-response.

For the stimulus-locked analysis, data from 200 ms before to 800 ms after the target were included, with baseline correction applied using the interval [-200, 0]. The response-locked analysis covered 800 ms before to 100 ms after the response, without baseline correction. Epochs containing artifacts were removed through semi-automated methods in EEGLAB, which identified improbable and abnormally distributed signals. Following EEGLAB’s pre-processing, event-related potentials were computed using ERPLAB.

#### ERP analysis

We examined the effect of noise level on two event-related potentials (ERPs), namely centro-parietal positivity (CPP) and lateralized readiness potential (LRP), through both stimulus-locked and response-locked analysis.

CPP, typically measured by averaging the amplitude of the electrodes CPz and its bilateral neighboring electrodes CP1 and CP2 (Kelly & O’Connell, 2013; 2015), was computed accordingly. In stimulus-locked analysis, a preliminary inspection of grand average ERP waveforms in those channels revealed the rise having approximately 350 ms onset. Hence, we analyzed [350 800] interval for stimulus-locked analysis. For response-locked analysis, we considered the [-450 0] interval to maintain consistent duration between both analyses. We employed one-way ANOVA on CPP amplitudes with the noise level as the main factor for both analyses. Pairwise comparisons were subsequently performed among the four noise levels. To explore further, we employed a more conservative approach utilizing false discovery rate (FDR) correction with the channels specified above and the time points included (Fields, 2017). For the stimulus-locked analysis, the time points from the stimulus onset to 800 ms post-stimulus; while for the response-locked analysis, the time points from 800 ms pre-response to the response onset were included. Scalp maps were also generated to visualize signal distribution across the entire brain during the epoch intervals. For LRP analysis, a measure of unimanual response preparation, a pair of electrodes from central and frontocentral regions in each hemisphere (C3-C4, and FC3-FC4) were included, in line with the literature (Eimer, 1998; Kelly & O’Connell, 2013). LRP for each trial was computed by subtracting the contralateral from ipsilateral channel activity with respect to the walking direction of the target stimuli in the trial. We initially utilized ANOVA with FDR correction including all time points following the stimulus or preceding the response onset. Following this, we also performed the analysis to include only FC3 and FC4 channels in the time interval approximately 100 ms later than the onset of CPP differentiation. Thus, in the latter analysis, the [450 800] interval for the stimulus-locked and the [-350 0] interval for the response-locked analysis were included. This analysis aimed to ascertain whether LRP temporally followed CPP, as suggested by Kelly and O’Connell (2013).

## RESULTS

### Behavior

The results of the behavioral analysis demonstrated that the mean response time and miss rate followed a trend aligned with the experiment’s difficulty level (see Figure 2). The mean response time (Figure 2, left) and miss rate (Figure 2, right) increased as the noise level increased. The mean response time for different noise levels ranges from 1 to 1.4 seconds, while the miss rate ranges from 1% to 25%.

**Figure 2.**
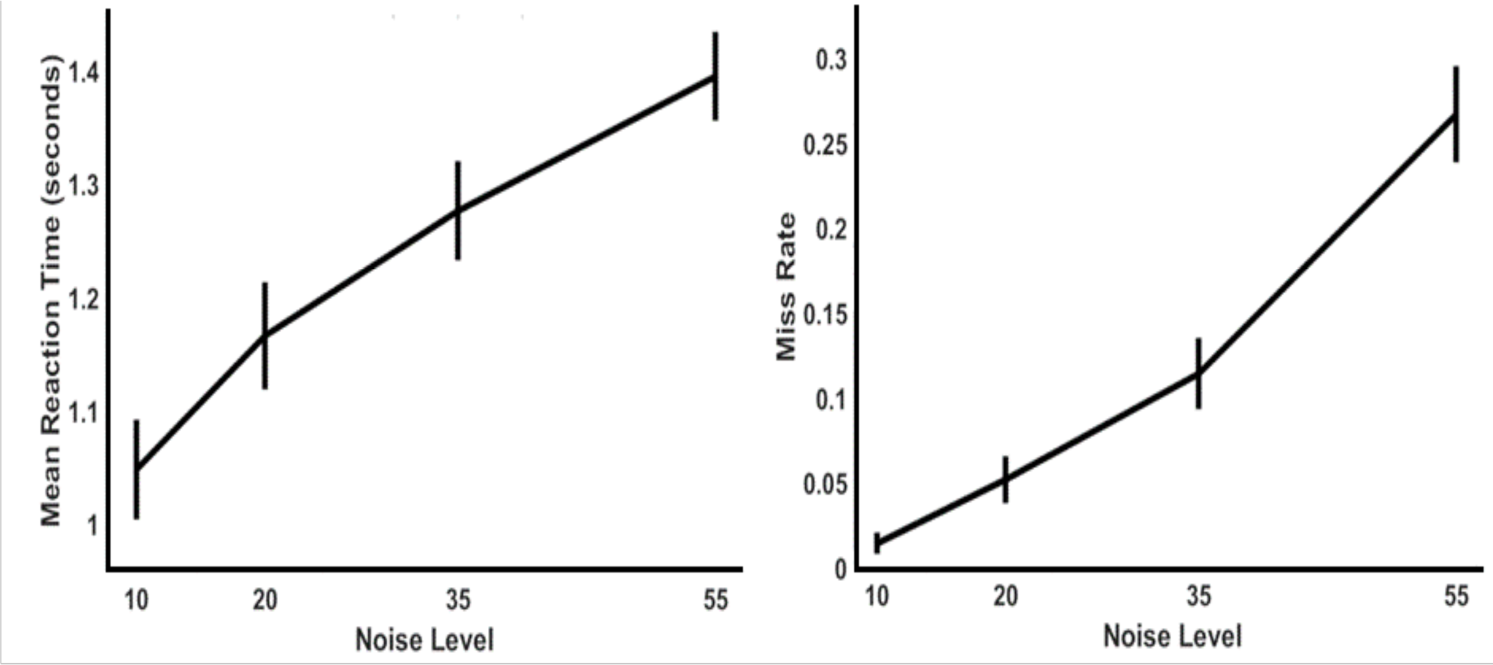
Behavioral Results. The mean reaction time (left) and the miss rate (right) are given according to the corresponding noise levels of the biological motion stimulus.

One-way ANOVA indicated a main effect of the noise level on the mean response time (F [3,45] = 246.953, p < 0.001, η_p_^2^=0.943), and all pairwise comparisons showed significant differences between the noise levels (p<0.001 for each comparison, see Table 1). Similarly, a main effect of the noise level was found on the miss rate (F [3,45] = 101.562, p<0.001, η_p_^2^= 0.871). Additionally, pair-wise comparisons between noise levels on miss rate revealed significant results (p<0.005), further providing support for the successful manipulation of difficulty in the experiment (see Table 1).

**Table 1.**
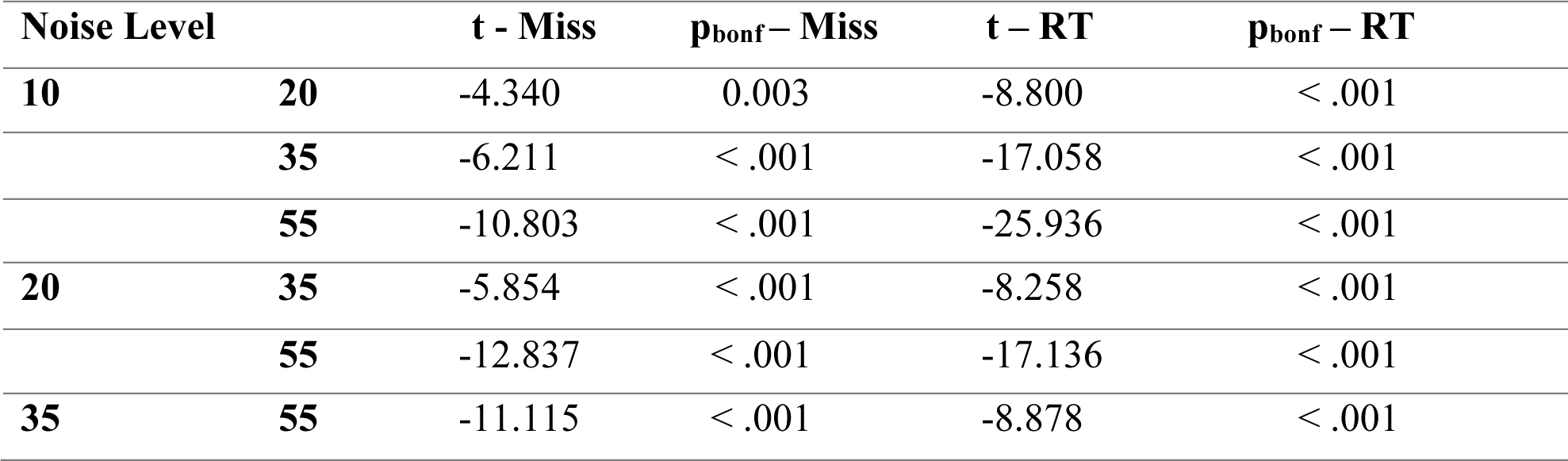
Pair-wise comparisons with corresponding t and p values for the miss rate (Miss) and mean reaction time (RT) across different noise levels.

| Noise Level |  | t - Miss | p<sub>bonf</sub> - Miss | t - RT | p<sub>bonf</sub> - RT |
| --- | --- | --- | --- | --- | --- |
| 10 | 20 | -4.340 | 0.003 | -8.800 | < .001 |
|  | 35 | -6.211 | < .001 | -17.058 | < .001 |
|  | 55 | -10.803 | < .001 | -25.936 | < .001 |
| 20 | 35 | -5.854 | < .001 | -8.258 | < .001 |
|  | 55 | -12.837 | < .001 | -17.136 | < .001 |
| 35 | 55 | -11.115 | < .001 | -8.878 | < .001 |

### Centro-Parietal Positivity (CPP)

#### Stimulus-Locked

To illustrate the effects of noise levels on CPP, we plotted the ERPs from stimulus onset to 800 ms post-stimulus. Complementing this, we generated scalp maps to visualize the general distribution of activity and its significance (See Figure 3A). Initial inspection of the plot revealed that CPP waveforms align with the difficulty level of the target stimuli, displaying lower peak amplitudes and slower build-up rates as noise levels increase. The scalp maps further underscored the increased activity for conditions with lower noise levels, with a distribution concentrated on the centro parietal brain areas.

**Figure 3.**
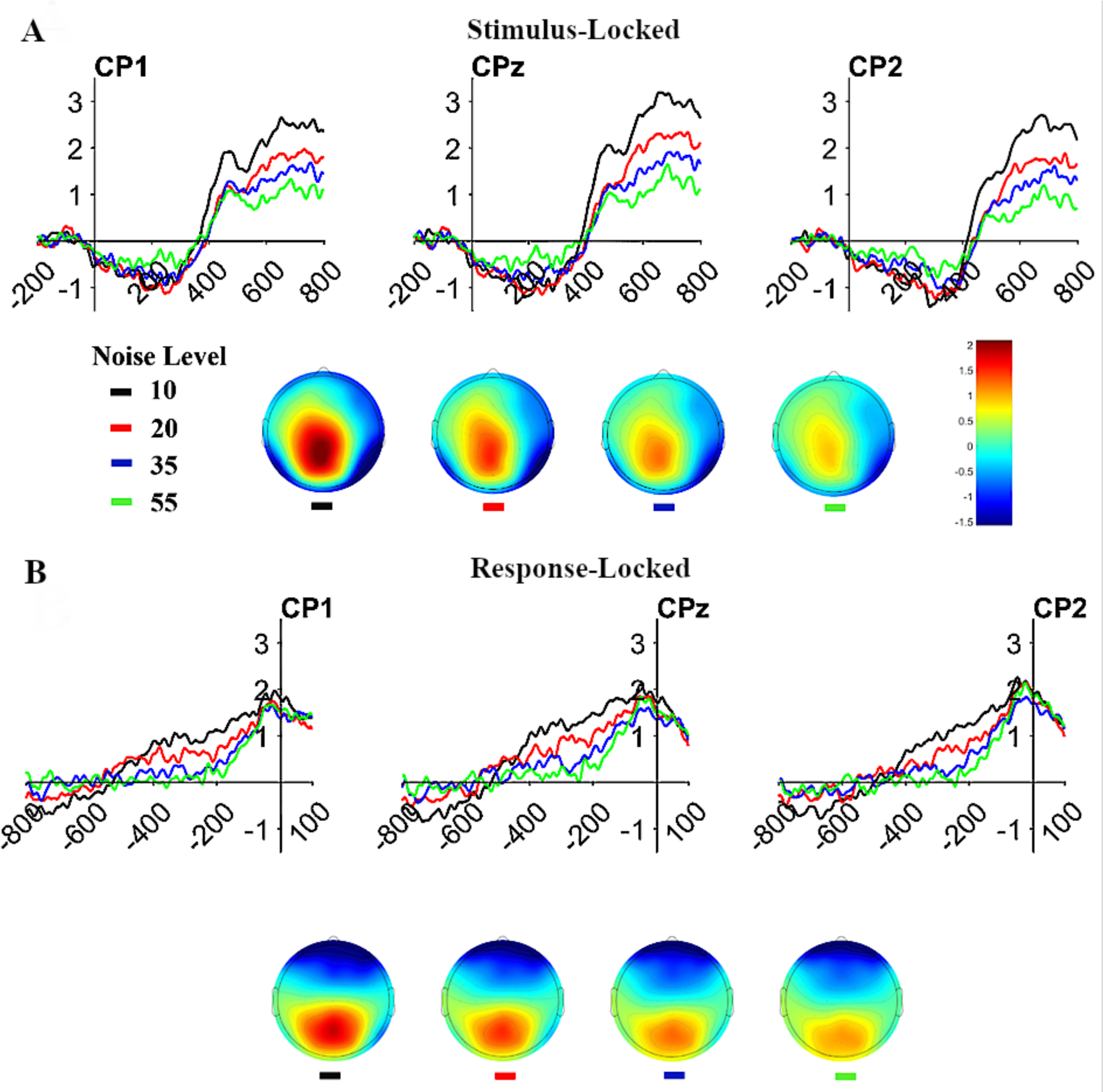
CPP results. Stimulus-locked (A) and response-locked (B) waveform graphs for CP1, CP2, and CPZ channels along with the scalp maps are shown.

Employing one-way repeated-measures ANOVA on mean amplitudes from CPP channels (CP1, CP2, and CPz) during [350 800], a notable main effect of noise level was found (F[3,6] = 196.615, p < .001, η_p_^2^=0.990). Following this, to investigate the effect of our manipulation more thoroughly, we performed pair-wise t-tests. All of the comparisons revealed significant effects after the Bonferroni correction (see Table 2).

**Table 2.** Pair-wise comparisons for the mean CPP amplitude in stimulus-locked analysis during [350 800] interval for different noise levels.

| Noise Level |  | Mean Difference | t | Cohen's d | p<sub>bonf</sub> |
| --- | --- | --- | --- | --- | --- |
| 10 | 20 | 0.617 | 13.228 | 2.346 | < .001 |
|  | 35 | 0.831 | 17.832 | 3.163 | < .001 |
|  | 55 | 1.081 | 23.187 | 4.113 | < .001 |
| 20 | 35 | 0.215 | 4.605 | 0.817 | 0.022 |
|  | 55 | 0.464 | 9.960 | 1.767 | < .001 |
| 35 | 55 | 0.250 | 5.355 | 0.950 | 0.010 |

In an exploratory effort to comprehensively assess the CPP activation across the entire time range, we employed ANOVA with FDR correction for the CPP channels and encompassing time points from the stimulus onset to 800 ms post-stimulus. In this analysis, the correction was done considering the number of channels and the time interval included to reduce the effect of false positives and hence Type I errors (Fields, 2017). This heightened level of conservatism enabled us to meticulously evaluate the significance of activations across CP1, CP2, and CPz, commencing right from the onset of the target stimulus. Remarkably, this analysis yielded a consistent pattern that mirrors the visual depiction of event-related potentials outlined above. Notably, the distinction in CPP responses based on varying noise levels began to emerge around 400 ms following the stimulus onset (see Supplementary Figure 1). This investigation further confirmed the chosen time interval in epoching.

#### Response-Locked

ERPs during [-800 100] for the response-locked analysis is illustrated along with the scalp maps (see Figure 3B). The figure highlights the alignment of CPP waveforms with the task difficulty, showcasing heightened peak amplitudes and build-up rates in lower noise level conditions. In addition, scalp maps indicate more robust activity in less difficult conditions. Noteworthy is the tendency for activity across noise levels to converge at a certain level during the response (at 0 point for the response-locked analysis).

Consistent with the stimulus-locked analysis, we employed first a repeated-measures ANOVA on the mean amplitudes during the [-450 0] epoch interval. This analysis revealed a main effect of noise level (F[3,6] = 276.680, p < .001, ηp2=0.993). Pair-wise tests between the noise levels also revealed significant differences between all comparisons except a marginally significant difference between noise levels 20 and 35 (See Table 3).

**Table 3.**
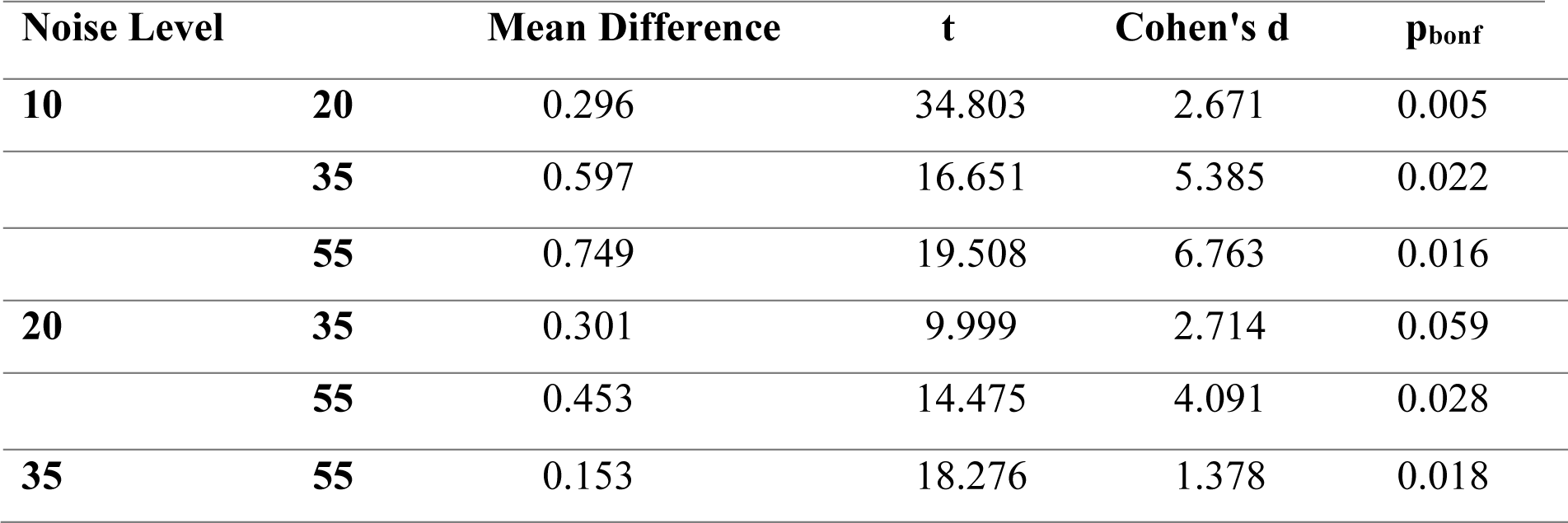
Pair-wise comparisons for the mean CPP activity in response-locked analysis during [-450 0] interval for different noise levels.

It is important to note that, as we have outlined above, the signals from each electrode showed a tendency to reach a certain level regardless of the difficulty level during response execution (See Figure 3B). This observation is also in line with the decision-making models that emphasize accumulation to bound dynamics. To further provide support for this trend, ANOVA with FDR correction was conducted on the [-800 0] time interval. The results of this analysis confirmed the trend with no main effect of noise on the activity observed in any of the CPP channels as the time approached the response onset (see Supplementary Figure 2).

### Lateralized Readiness Potential (LRP)

#### Stimulus-Locked

To measure the lateralized readiness potential (LRP), we calculated the difference between contralateral and ipsilateral channels based on the target’s walking direction. We included the electrodes from central and fronto-central sites as pairs (C3 and C4, FC3 and FC4) in line with the previous literature (Kelly and O’Connell, 2013). The LRP waveforms obtained from each pair during the epoched time interval for the stimulus-locked analysis were plotted in Figure 4A. Employing ANOVA with FDR correction analysis on the mean activity during the period from the stimulus onset to 800 ms post-stimulus, we found that LRP activity did not exhibit the same alignment with task difficulty observed in CPP activity.

**Figure 4.**
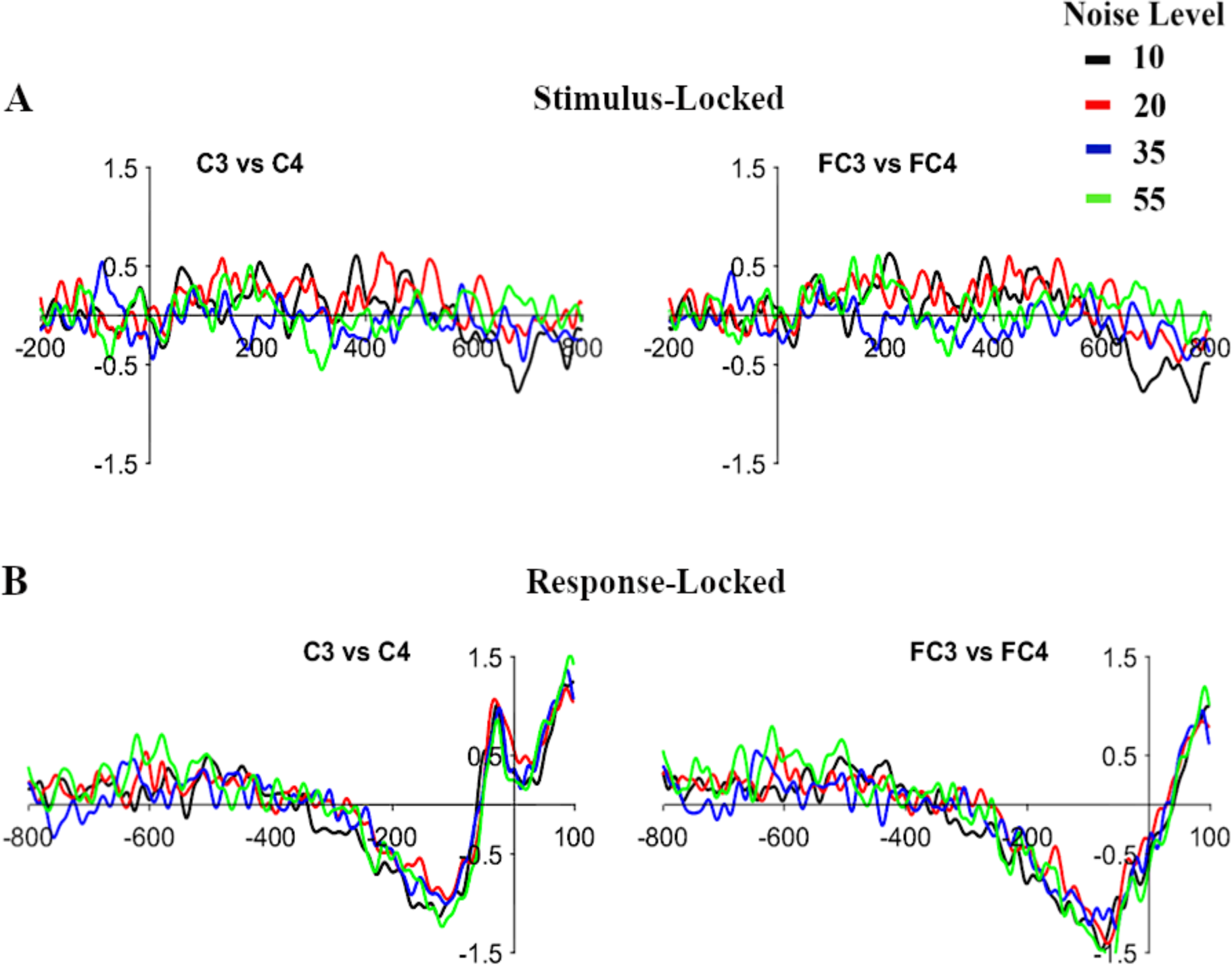
LRP Results. Stimulus-locked (A) and response-locked (B) LRP activities are shown for two different pairs of electrodes, namely C3 – C4 and FC3 – FC4.

Following this, we performed a modified version of this analysis akin to the similar study done by Kelly & O’Connell (2013), where they detected a delayed LRP activity measured in frontocentral sites for approximately 100 to 150 ms in relation to CPP formation. Thus, we focused our analysis on the mean activity during the interval from 450 ms to 800 ms post-target, excluding C3 and C4 electrodes and narrowing it down to FC3 and FC4. Nevertheless, this analysis also failed to demonstrate a significant main effect of the noise level in the stimulus-locked data.

#### Response-Locked

Similar to the stimulus-locked analysis, we plotted LRP waveforms for each electrode pair within the response-locked epoch (see Figure 4B). Applying ANOVA with FDR correction analysis by considering the mean activity during the time interval from 800 ms pre-response to response onset, the alignment with the difficulty level was not observed as in the stimulus-locked analysis.

Interestingly, further modified analysis conducted on the response-locked data considering the mean activity during the period from 350 ms pre-response to response onset demonstrated a significant main effect of noise level, providing support for generalizing the previous findings in the literature (F[3,45] = 3.3468, p = .03). The discrepancy between stimulus-locked and response-locked results likely arises because of the stimulus-locked analysis not being well-suited to capture the response preparation interval, as compared to more insightful response-locked analysis.

## DISCUSSION

The present study aimed to investigate the neural dynamics of biological motion perception in the framework of perceptual decision-making, thereby integrating two lines of research. To achieve this, the experiment employed PLDs with varying degrees of noise dots to manipulate the difficulty level in the point-light walker direction discrimination task.

The behavioral results highlighted that overall task performance improved notably under conditions of lower noise. In essence, biological motion stimuli with heightened noise levels translated into delayed responses and an increased rate of misses, verifying our manipulation of task difficulty. On the other hand, investigating the electrophysiological results provided a more comprehensive overview of the neural underpinnings. Specifically, the amplitude of the CPP exhibited an upward trajectory and earlier peak under lower noise conditions. Furthermore, signal distribution was consistent with the previous findings, emphasizing heightened activation in the central brain sites corresponding to the CP1, CP2, and CPz channels, which are the primary focus for CPP. Moreover, the activity in those channels exhibited a pattern of reaching a common level during response irrespective of the task difficulty. These findings bolstered the essential qualities of CPP, highlighting its role in indexing sensory evidence accumulation over time until a certain threshold is attained during the response. Another important implication of the results is the differentiation of motor preparation signal LRP from CPP. It has been found that LRP lagged the CPP formation for approximately 100 to 150ms. In sum, this study’s outcomes resonated with the prior literature, solidifying evidence for a domain-general mechanism driving the perceptual decision-making process.

### Impact on the biological motion perception literature

Biological motion perception has been extensively studied in the literature, predominantly employing point-light displays. Utilizing non-invasive techniques like fMRI and EEG, previous studies have identified relevant brain areas including posterior superior temporal sulcus (Vaina et al., 2001), and explored the computational aspects through modeling (Giese & Poggio, 2003; Lange & Lappe, 2006; Duarte et al., 2022). Event-related potential (ERP) studies have primarily focused on form, motion, and configuration processing, with a particular emphasis on early ERP components like N1, P1, N200, and N330 (Hirai et al., 2005; Baccus et al., 2009). Additionally, some studies have delved into higher-level processes involving attention, visual search and action recognition (White et al., 2014; Hirai & Hiraki, 2006). Notably, a few influential studies have identified a centro-parietal deflection resembling CPP at around 300ms, suggesting potential involvement in later cognitive processes (Krakowski et al., 2011). To the best of our knowledge, the present study fills an important research gap by explicitly investigating perceptual decision-making and related ERPs in the context of biological motion perception, introducing a novel framework and a new avenue to pursue in the field.

Besides being the potential benchmark for inspiring further empirical studies, the present study also holds the potential to improve computational models of biological motion perception. One of the most influential models was developed by Giese and Poggio (2003). This model suggests that biological actions are performed in two streams, namely the ventral and dorsal streams, that process form and motion, respectively. According to this model, convergence at the superior temporal sulcus integrates these pathways to yield the perception of motion (Giese & Poggio, 2003). However, this model has limitations since it relies on solely feed-forward mechanisms without incorporating the top-down processes. Notably, the concurrent processing of decisions within the two streams or after form-motion integration remains to be investigated. Therefore, the integration of perceptual decision-making in the framework of this model would provide further insights into the underlying mechanisms of biological motion perception. Other models have also been proposed to explain the global structure-from-motion, local motion processing stages, and infant perceptual abilities (Lange & Lappe, 2006; Hirai & Senju, 2020; Troje, 2008). Similarly, these models also stand to benefit from our endeavor, in terms of enhancing comprehensiveness and accuracy.

### Impact on the perceptual decision-making literature

The present study not only replicates prior research on perceptual decision-making but also introduces a more intricate task involving motion clips, setting it apart from studies that utilize simpler or static stimuli like face versus car discrimination or random-dot motion tasks. (Philiastides & Sajda, 2005; Newsome et al., 1989). Consequently, the present study extends previous research by employing a more ecologically valid and socially relevant task. Moreover, the results revealed a noteworthy delay in CPP formation compared to previous studies, highlighting the responsiveness of perceptual decision-making and CPP to stimulus complexity.

In terms of implications, the domain-general mechanism of perceptual decision-making can manifest through distinct networks. Notably, ancillary circuits, that play a complementary role to core decision circuits, may exhibit variations depending on the specific task and stimuli employed (O’Connell & Kelly, 2021). The interactions between stimulus-specific regions that lead to appropriate responses deserve further scrutiny. Specifically, the role of pSTS merits deeper exploration using a similar paradigm, as this region has been also linked with biological motion perception (Saygin et al., 2004), and is considered to be a convergence point for dorsal and ventral pathways (Giese & Poggio, 2003). Additionally, studying the network connecting pSTS and homologous parietal regions in humans to macaque LIP holds promise, akin to research exploring differences between the neural responses of MT and LIP regions (Gold & Shadlen, 2007).

Concurrently, models of perceptual decision-making can benefit from socially meaningful and ecologically valid stimuli, such as biological motion. Investigating whether the drift rate of sensory accumulation or the decision-boundary is influenced by stimulus quality that leads to later rise in CPP in the present study is an avenue worth exploring. The recent models incorporate concepts such as drift rate, speed-accuracy trade-offs, urgency signals, and uncertainty signals (O’Connell & Kelly, 2021; Hanks & Summerfield, 2017). Likewise, studying processes like accuracy-speed trade-offs, urgency, or uncertainty signals gains significance when considering biological motion stimuli, given its potential to reflect evolutionarily meaningful scenarios.

### Limitations and future research

The present study bears its first limitation concerning the stimuli used. Although temporal qualities of scrambled motion are manipulated by varying the display time, there are also other potential manipulations. Importantly, it would be worth exploring the manipulation of the speed of the stimuli used in the task. Such an investigation could alter the CPP onset and latency, revealing potential nonlinear effects of speed on the sensory evidence accumulation process. In addition, point-light displays, while effective in capturing crucial aspects of biological motion, could be complemented by more naturalistic videos depicting human actions or even real human actors in naturalistic settings. This avenue of research aligns with the growing adoption of real-time EEG methodologies, offering the opportunity to enhance ecological validity (Stangl et al., 2023; Pekçetin et al., 2023).

It should also be noted that further research is needed to investigate whether CPP reflects the two essential components of the drift-diffusion model. Recent endeavors in computational modeling, notably neurally informed models, extend beyond traditional drift-diffusion models while explaining more complex mechanisms. With the inclusion of time constraints and speed-accuracy trade-offs, these models offer a promising direction for future investigations (Kelly et al., 2021).

One should also note that while the task employed in the present study focused on visual perception, the integration of sensory information holds great significance in our daily experiences. This holds true for biological motion perception in natural settings since it consists of and requires multisensory input (Brooks et al., 2007). Despite the prevailing use of visual motion displays, the exploration of audio-visual cues for human motion in natural settings is a growing area of research (Mendonça et al., 2011). Therefore, it would be worth investigating the effects of multisensory integration on the CPP build-up using a similar experiment. Anticipated benefits of multimodal integration include more efficient sensory evidence accumulation which may potentially lead to earlier latencies and heightened peaks in CPP.

## Conclusion

In conclusion, the study advances our understanding of perceptual decision-making by investigating its dynamics within the context of biological motion perception, offering a fresh perspective that can guide future investigations. The framework proposed in the present study is promising for building more comprehensive computational models and innovative research endeavors for both fields.

## SUPPLEMENTARY FIGURES

**Supplementary Figure 1:**
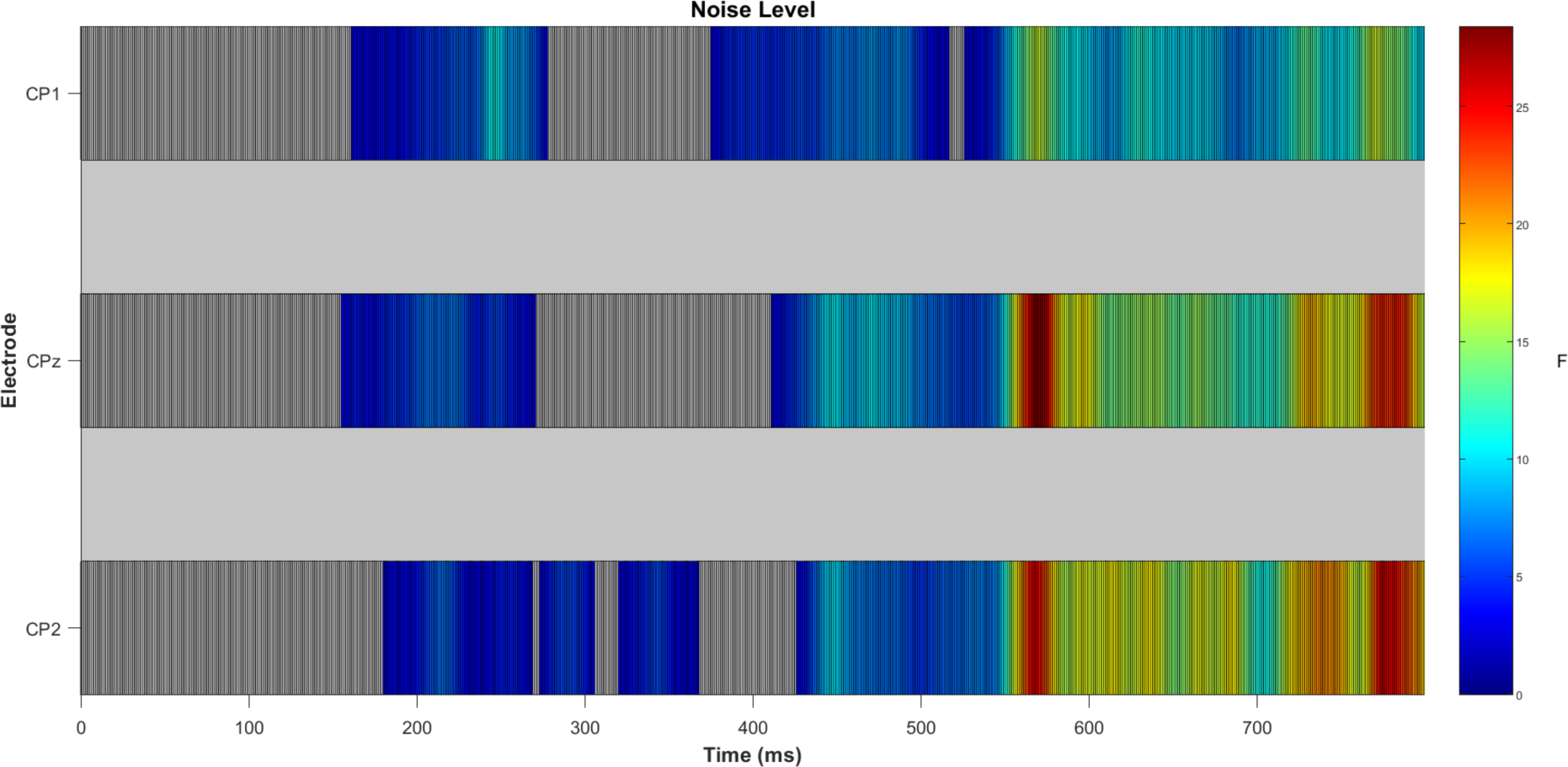
Activity in CPP channels starting from the stimulus onset to 800 ms post stimulus.

**Supplementary Figure 2:**
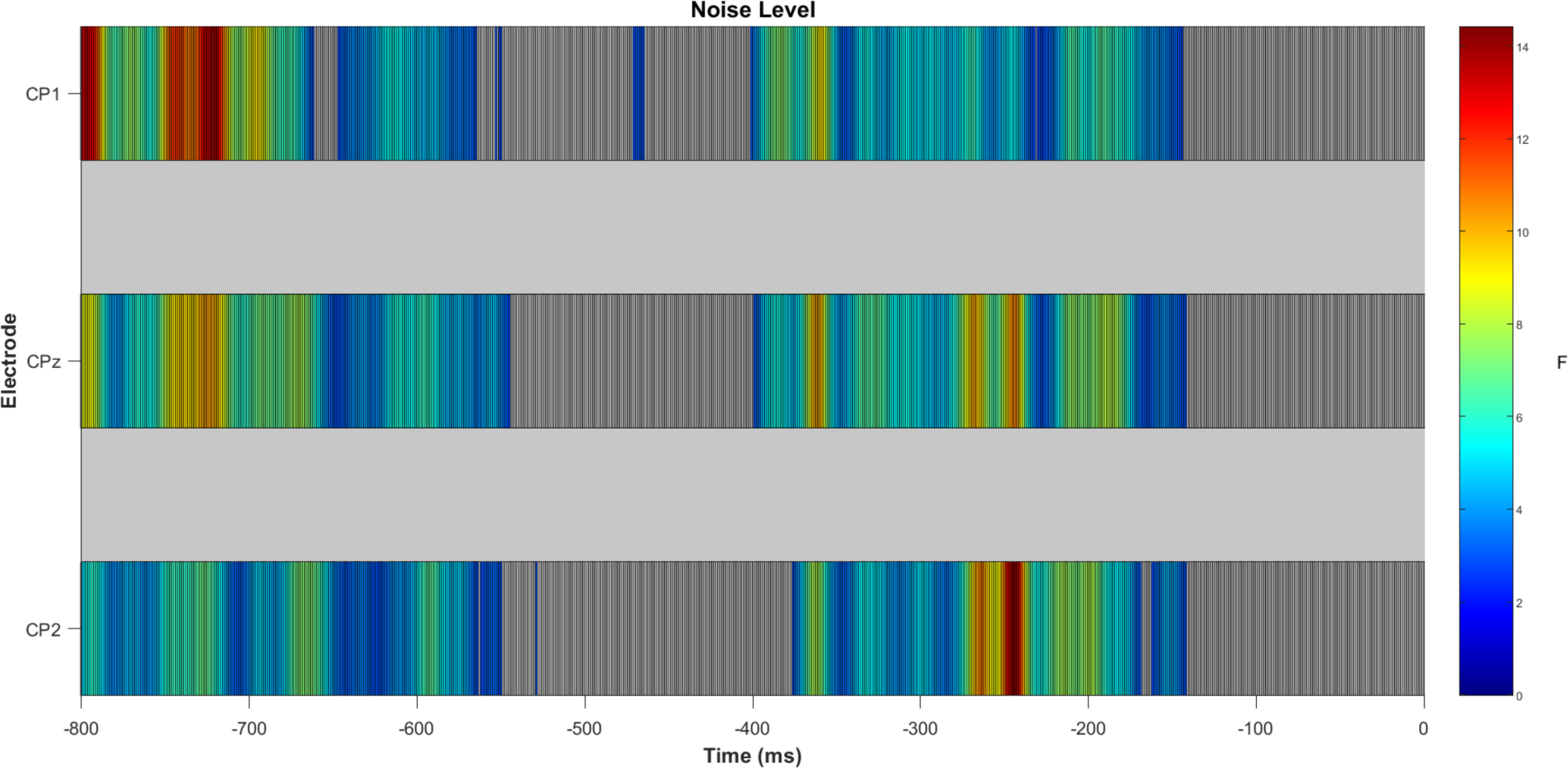
Activity in CPP channels starting from 800 ms pre-response to the response onset.

